# TaxaHFE: A machine learning approach to collapse microbiome datasets using taxonomic structure

**DOI:** 10.1101/2023.06.06.543755

**Authors:** Andrew Oliver, Matthew Kay, Danielle G. Lemay

## Abstract

**Motivation:** Biologists increasingly turn to machine learning models not just to predict, but to explain. Feature reduction is a common approach to improve both performance and interpretability of models. However, some biological data sets, such as microbiome data, are inherently organized in a taxonomy, but these hierarchical relationships are not leveraged during feature reduction. We sought to design a feature engineering algorithm to exploit relationships in hierarchically organized biological data.

**Results:** We designed an algorithm, called TaxaHFE, to collapse information-poor features into their higher taxonomic levels. We applied TaxaHFE to six previously published datasets and found, on average, a 90% reduction in the number of features (s.d = 5.1%) compared to using the most complete taxonomy. Using machine learning to compare the most resolved taxonomic level (i.e., species) against TaxaHFE-preprocessed features, models based on TaxaHFE features achieved an average increase of 3.47% in receiver operator curve area under the curve (ROC-AUC). Compared to other hierarchical feature engineering implementations, TaxaHFE introduces the novel ability to consider both categorical and continuous response variables to inform the feature set collapse. Importantly, we find TaxaHFE’s ability to reduce hierarchically organized features to a more information-rich subset increases the interpretability of models.

**Availability and Implementation:** TaxaHFE is available as a Docker image and as R code at https://github.com/aoliver44/taxaHFE.

## INTRODUCTION

With the cost of DNA sequencing continuing to drop faster than compute power increases (Wetterstrand) the analysis of large data sets remains a bottleneck in biological research. One method for analysis is machine learning (ML) (Choi *et al*., 2022), which is a blanket term that refers to computer algorithms designed to find patterns in data, iteratively optimizing performance without human input. While ML methods represent a suite of powerful and sensitive tools, they can suffer from a problem present in many human -omic studies: many features (i.e., microbial taxa) describing relatively few samples. Mathematician Richard Bellman referred to this problem as the “curse of dimensionality” (Bellman, 2003). Practically speaking, too many features can result in “overfitting”, leading to poor generalizability of the model. For this reason, implementing methods to reduce the size of data while retaining its important features can improve both the speed, the generalizability, and interpretability of the data.

Feature engineering, or the set of preprocessing steps done to data prior to ML model evaluation, can help address problems imposed by high-dimensional data. While illustrating the totality of feature engineering is beyond the scope of this paper, some general examples include scaling or normalizing features, removing low variance features, collapsing highly correlated features, sophisticated methods for selecting subsets of features, and collapsing features into principal coordinate space. The goal of several of these methods is to reduce the feature space (dimensionality) and produce a highly discriminatory set of variables with respect to a response of interest. However, in biology models are not just used to make predictions, but to *explain*. This means that the reduced feature set needs to also be interpretable, rather than alternate ordinations, such as principal components.

Some biological data, such as microbiome and dietary data, can be represented using hierarchical structures (Jacobs and Steffen, 2003; Johnson *et al*., 2019; Choi *et al*., 2022). Taxonomic assignments have long been used to identify microorganisms. This taxonomy is usually represented by a hierarchical classification scheme, whereby the ancestral level is the most general group, followed by increasingly specific grouping rules. More recently, researchers have begun to represent consumed foods in a similar taxonomic way (Johnson *et al*., 2019). More than merely identifying information, taxonomy represents relatedness, reflecting ecological patterns (for microorganisms) (Bevilacqua *et al*., 2021) or complex admixtures of similar chemicals and nutrients (for food) (Johnson *et al*., 2019). While data with hierarchical structure presents many different levels by which to analyze a trait or response, researchers generally choose to collapse these data to a single level for ease of analysis (i.e., analyzing microbiome data at the family-level) (Kleine Bardenhorst *et al*., 2021). This can be a useful strategy, especially if the trait/response of interest is known to be conserved at a certain phylogenetic depth which can be approximated by a taxonomic level (Martiny *et al*., 2015). Even without knowledge of a conserved phylogenetic depth *a priori*, tools exist to identify the average phylogenetic depth of a trait/response; however, many of these tools require a phylogenetic tree as input (Martiny *et al*., 2012). Often there is not *a priori* information available, and summarizing taxa to a specific level is weakly justified. Even more, if the data was summarized at a taxonomic level, but the response to a treatment is conserved at a different taxonomic level, it could lead to a false conclusion that, for example, the microbiome does not respond to a treatment of interest. A systematic review of current practices for the analysis of human microbiome data revealed a lack of consensus about which taxonomic level to study (Kleine Bardenhorst *et al*., 2021). Why not let information theory decide?

Here we introduce a method for hierarchical feature engineering (HFE) to dynamically collapse hierarchical data purely based on taxonomic relationships and information gain. Our algorithm, TaxaHFE, does not require the user to have knowledge *a priori* regarding the taxonomic level of conservation for a given trait. Rather, it seeks to maximize the information contained at various taxonomic levels while simultaneously reducing redundancy in the feature space. As a proof of concept, we apply TaxaHFE to microbiome data and compare it to an existing hierarchical feature engineering algorithm (Oudah and Henschel, 2018), assessing feature reduction and downstream ML performance. Additionally, we show that TaxaHFE’s utility extends to other hierarchically organized data by applying our algorithm to hierarchical food data represented by taxonomic trees.

## METHODS

### TaxaHFE algorithm

The algorithm (Figure 1) can broadly be broken down into two main sections: 1) the creation of a taxonomic tree representing the hierarchical data, and 2) the competitions of each taxon in a post-order tree traversal. Within the competition section, four major steps occur: 1) a feature abundance and prevalence filter, 2) a correlation competition between parent and child taxa, 3) a machine learning step to determine information content of the taxa from the previous step, and 4) one additional machine learning step on all the “winning” features. These steps are graphically represented in Figure 1 and algorithmically outlined below.

**Figure 1:**
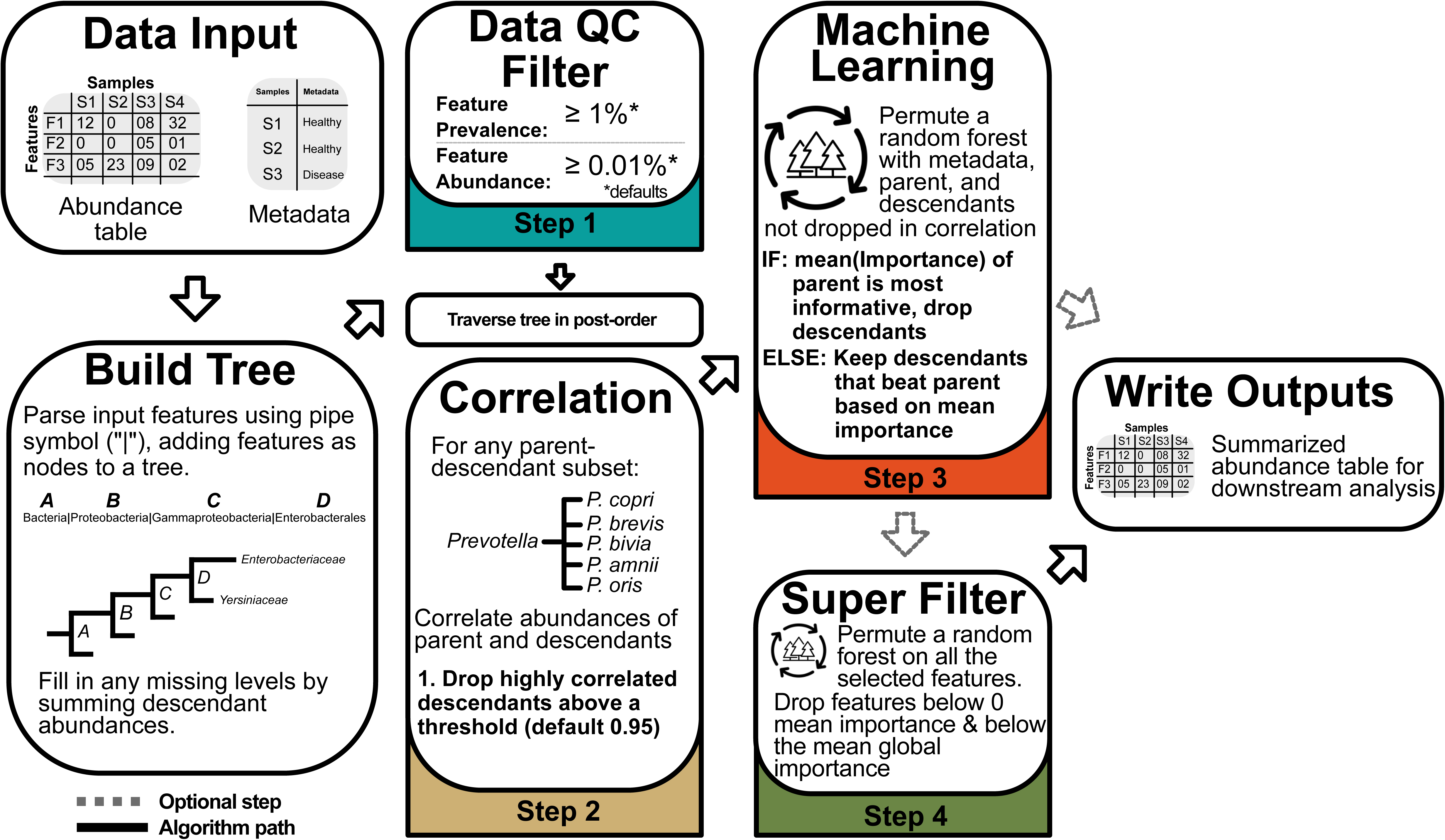
Overview of TaxaHFE

#### 1. Build tree

a. Generate a node *x* in the tree **T** for each taxon
b. Store the abundance values for the taxon on *x*, as well as calculating whether the abundances meet minimum mean abundance and prevalence thresholds (Figure 1, Step 1)
c. To fill in any missing abundance data, traverse the tree **T** in post-order, generating a missing abundance vector *a_n_* for a node *x_n_* as the vector sum of all abundance vectors from the direct descendants *c_n_ = children(x_n_)*, where *a_c_* is the abundance vector of child node *c*.

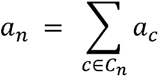

#### 2. Compete tree

Traverse the tree **T** in post-order, considering every subtree **T***_n_* using the following steps.

a. If *root*(**T***_n_*) has not met the minimum mean abundance and prevalence thresholds, this taxon will not be considered further (Figure 1, Step 1)
b. If *root*(**T***_n_*) is a leaf node, mark “winner” and proceed to the next subtree

i. Note: Being marked a “winner” is temporary, and any “winner” in a particular subtree ***T_n_*** at level *l* will be reconsidered at level *l* − 1 during the traversal.
c. Traverse ***T_n_***, generating a set **N_0_** of nodes previously marked as “winner”. If a node *x* is marked “winner”, no descendent nodes are considered.
d. Generate a subset **N_nc_** ∈ **N_0_**, containing each node *x* in **N_0_** whose abundance vector *a_x_* is not correlated with the abundance vector *a_n_* of *root*(**T***n*), above the specified threshold t (Figure 1, Step 2). For all nodes **N_0_** - **N_nc_**, remove the “winner” designation:

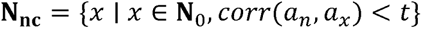
e. Using the set of abundance vectors **A**= {*a_n_, a_nc_*_1_, *a_nc_*_2_,…) (from *root*(**T***_n_*) and the non-correlated nodes **N_nc_**) and response variables **M** (from the metadata input), fit a random forest model to determine the taxa importances to **M** calculated using Gini impurity-corrected scores (Nembrini *et al*., 2018) and generating a vector of scores **S** (Figure 1, Step 3).

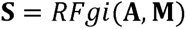 Any node *x* with a score *s_x_* greater than *root*(**T***_n_*) score *s_n_* is marked “winner”

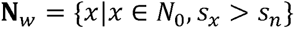 Otherwise, if the score of ***root***(**T*_n_***) is the highest value, only it is selected

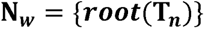 All other nodes **N_nc_** - **N_*w*_** have the “winner” designation removed, as well as ***root***(**T***_n_*) if ***root***(**T***_n_*) ∈ N*_w_*. **Traversal will be stopped at a level *l***, where *l* ≥ 2, such that ***level***(***root***(**T**_n_)) ≥ ***l*** for all competed subtrees **T**_c_. This prevents ***root***(**T**_0_) (that contains the sum of all abundance vectors) from being included in the competition, and also allows for the preservation of taxonomic information at a definable level. Because they have not been traversed, any node ***x*** where ***level***(***x***) < l is not considered in the algorithm and cannot be marked “winner”. The result of this is a set of distinct nodes **N*_w_***, marked as “winner” across all competed subtrees **T_*c*_** having ***root***(**T_0_**) (the root of the full tree) as the only common ancestor. An optional final random forest model (Figure 1, Step 4) is then fit using the set of abundance vectors **A_*w*_** from **N_*w*_**, and the response variables **M**.

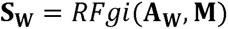 The final feature set **N_f_** is the set of nodes ***x*** with scores ***s_x_*** greater than zero and the average score Gini impurity-corrected scores of **N_f_**:

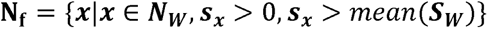

### Input data

The input data to TaxaHFE is (1) a flat file (comma- or tab-delimited) with a column labeled “clade_name” containing the taxonomic features with subsequent columns as samples containing numeric feature abundances and (2) a response variable flat file (Figure 1). The levels of the features within the input data should be delineated by a pipe symbol (“|”). For example, if the input data are microbiome features, the features should be Kingdom|Phylum|Class|…|last level. The taxonomic features can contain any number of levels present. The sample column names must be unique for each column in the input file and match a column in the response variable file, which also must contain a column encoding the response of interest (either categorical or continuous).

### Implementation

A reference implementation of this algorithm was written in R as part of development. The implementation uses the data.tree package (v1.0.0) ( Glur *et al*., 2020) to represent the hierarchical structure of input taxa, as well as for the post-order traversal of the data used in several places. Data.tree parses the input file column by using the pipe symbol (“|”) found in the input data, described above. Once imported into TaxaHFE, four broad steps occur, as described in the algorithm above: 1) feature abundance and prevalence filters, 2) a correlation competition between parent taxon and child taxa, 3) a random forest competition selecting the most informative (relative to the response variable) features from the previous step and 4) and an optional final “super filter” (SF) random forest of all the taxa that have survived from steps 1-3 (Figure 1). TaxaHFE applies reasonable defaults to these steps. For instance, by default, features are only considered if they have a minimum mean feature abundance of 0.01% and a minimum feature prevalence of 1%. Both filters can be set to zero or any other number, depending on the scale of the input data. Additionally, if the user is predicting a rare class (i.e., class imbalance), if may be prudent to keep rare features by setting these filters to zero. For the correlation competition, a child taxon abundance must be correlated with the parent abundance at less than 0.95 Pearson correlation to pass to step three. Finally, both step three and step four use random forests (RF) to report back Gini impurity corrected scores per feature, indicating the feature’s ability to predict the user-supplied response variable. These RF models are built and rebuilt 40 times by default, averaging out the Gini impurity scores per feature. The RF model fitting is done using the ranger package (v0.14.1) (Wright and Ziegler, 2017), including the ability to distinguish between numeric and factor response variable types in the input metadata. Randomness is seeded in the implementation using system time, but an allowance is made for defining a specific random seed. This randomness influences the outcome of the RF competitions run by the ranger package. For ease of use, the implementation has been released on GitHub and additionally packaged into a Docker container, allowing it to be run reliably in a variety of computing environments.

### Output data

The outputs of TaxaHFE are individual files of summarized original abundances for each hierarchical level and two files containing either the TaxaHFE or the TaxaHFE + SF selected features alongside the response variable.

### Downstream machine learning

The output of TaxaHFE is a reduced feature set, which may increase the performance of predictive analyses such as machine learning. To test this, we analyzed hierarchical datasets using a machine learning pipeline based around the Tidymodels package (v1.0.0) (Kuhn and Wickham, 2020) in R. Input data was split using 70% data for training and hyperparameter tuning, and 30% was used for testing. Inside the training split, 10-fold repeated (3x) cross validation was used for Bayesian-search hyperparameter tuning. A gentle correlation feature reduction step was also introduced within each cross-validation fold, yet at 0.95 Pearson (the same as TaxaHFE’s internal correlation filter), it likely had minimal effect on ML performance. The search space was limited to 80 different hyperparameter combinations (optimizing mtry and minimum node size) or ten minutes of search time, whichever finished first. Bayesian hyperparameter search was allowed to preemptively end if the scoring metric was not improved after ten iterations. The metric optimized was balanced accuracy for classification or mean absolute error for regression. The best model was fit to the left-out test data and assessed using area under the receiver operator curve (ROC-AUC), balanced accuracy, and Cohen’s kappa (Kuhn *et al*., 2023). For multi-class models, the Hand-Till ROC-AUC (Hand and Till, 2001) and macro-averaged balanced accuracy was used. Each ML model was run using ten different random seeds to reduce stochasticity introduced from different train-test splits. To determine the importance of features, the package fastshap (v0.0.7) (Greenwell, 2021; Štrumbelj and Kononenko, 2014) was used. Briefly, the best model was fit to the entire input data, and Shapley values were calculated for each feature. The R package ShapViz (v0.4.1) (Mayer and Stando, 2023) was used to plot these values.

### Evaluation of TaxaHFE

Six previously published microbiome datasets (Lloyd-Price *et al*., 2019; Mars *et al*., 2020; Muller *et al*., 2022; Wang *et al*., 2020; Franzosa *et al*., 2019; Erawijantari *et al*., 2020; Oliver *et al*., 2022) and a dietary tree dataset (Kable *et al*., 2022) were used to assess TaxaHFE (Supplemental Table 1). For the microbiome datasets, TaxaHFE parameters were set to minimum mean feature abundance of 0.01% and minimum feature prevalence of 1%. The correlation threshold was set to 0.95, and nperm = 40 (defaults for TaxaHFE) and the random seed was set to 42. For the food dataset, the abundance filter was changed to 0. Each dataset was summarized at either the order, family, genus, or species level (using TaxaHFE, which applies the prevalence and abundance filters prior to writing the summary files). Additionally, the species summarized microbiome data was analyzed using a previously published algorithm (hfe_algorithm.py (Oudah and Henschel, 2018)), using two correlation cutoffs: the program’s default 0.7 and 0.95 (matching TaxaHFE’s default correlation filter). These summarized and TaxaHFE-selected data were used in machine learning models to predict response variables associated with each study (Supplemental Table 1), as described above. To test whether ML performance was significantly different across different feature reduction methods, we used a linear mixed-effects model (from the R package nlme v3.1-157 (Pinheiro *et al*., 2019)), with study*level as a fixed effect and the random seed as a random effect. Quantile-quantile plots were investigated for normality of residuals. Study-specific models were also built like the above model, without the study interaction term. Finally, we performed estimated marginal means (EMMs) post-hoc tests using the emmeans package (v1.8.8) (Lenth *et al*., 2018) with Bonferroni adjusted p-values. We also analyzed the features compositionally by measuring the variance explained by the features selected using PERMANOVA models. To do so, we used the adonis(method = “bray”, nperm = 999) function from the vegan (v2.6-4) package (Oksanen *et al*., 2019) in R. For comparative purposes, Lefse (Segata *et al*., 2011) was run on Galaxy (http://galaxy.biobakery.org/) using default parameters, and Boruta was run using the Boruta package (v8.0.0) (Kursa and Rudnicki, 2010) in R using default parameters.

### Software and data availability

The code for TaxaHFE, along with installation instructions and example inputs, can be found on GitHub (https://github.com/aoliver44/taxaHFE). TaxaHFE version 2.0 was used for all analyses. The ML pipeline to assess the performance of TaxaHFE can also be found on GitHub (https://github.com/aoliver44/nutrition_tools). Location of datasets used for comparisons can be found in Supplemental Table 1. We downloaded five of the microbiome datasets from a microbiome-metabolome dataset collection (Muller *et al*., 2022).

## RESULTS

### TaxaHFE improves feature reduction compared to alternative taxonomically informed methods

We initially investigated how well TaxaHFE performs compared to summarizing microbiome datasets to higher taxonomic levels (order (L4) to species (L7)), or when using a previously published hierarchical feature engineering program (Oudah_70 and Oudah_95). In all six previously published studies, the dimensional reduction produced by TaxaHFE (± super filter, SF) selects features which, when used as input in machine learning models, result in the higher mean ROC-AUC scores compared to data summarized at specific taxonomic levels, or a previously published HFE algorithm (Figure 2A, Supplemental Figure 1, Supplemental Table 2). The mean ROC-AUC for TaxaHFE (+SF) across all studies assessed was 0.901 (s.d. = 0.071) followed by TaxaHFE (-SF) (0.897, s.d. = 0.071). Moreover, TaxaHFE (+SF) produces the best models for 4/6 studies when assessed using balanced accuracy (Figure 2B, Supplemental Table 2) and 4/6 studies using mean Cohen’s kappa (Figure 2C, Supplemental Table 2). In the other two cases, TaxaHFE preprocessed models were not significantly different from the best models produced (Supplemental Figure 1).

**Figure 2:**
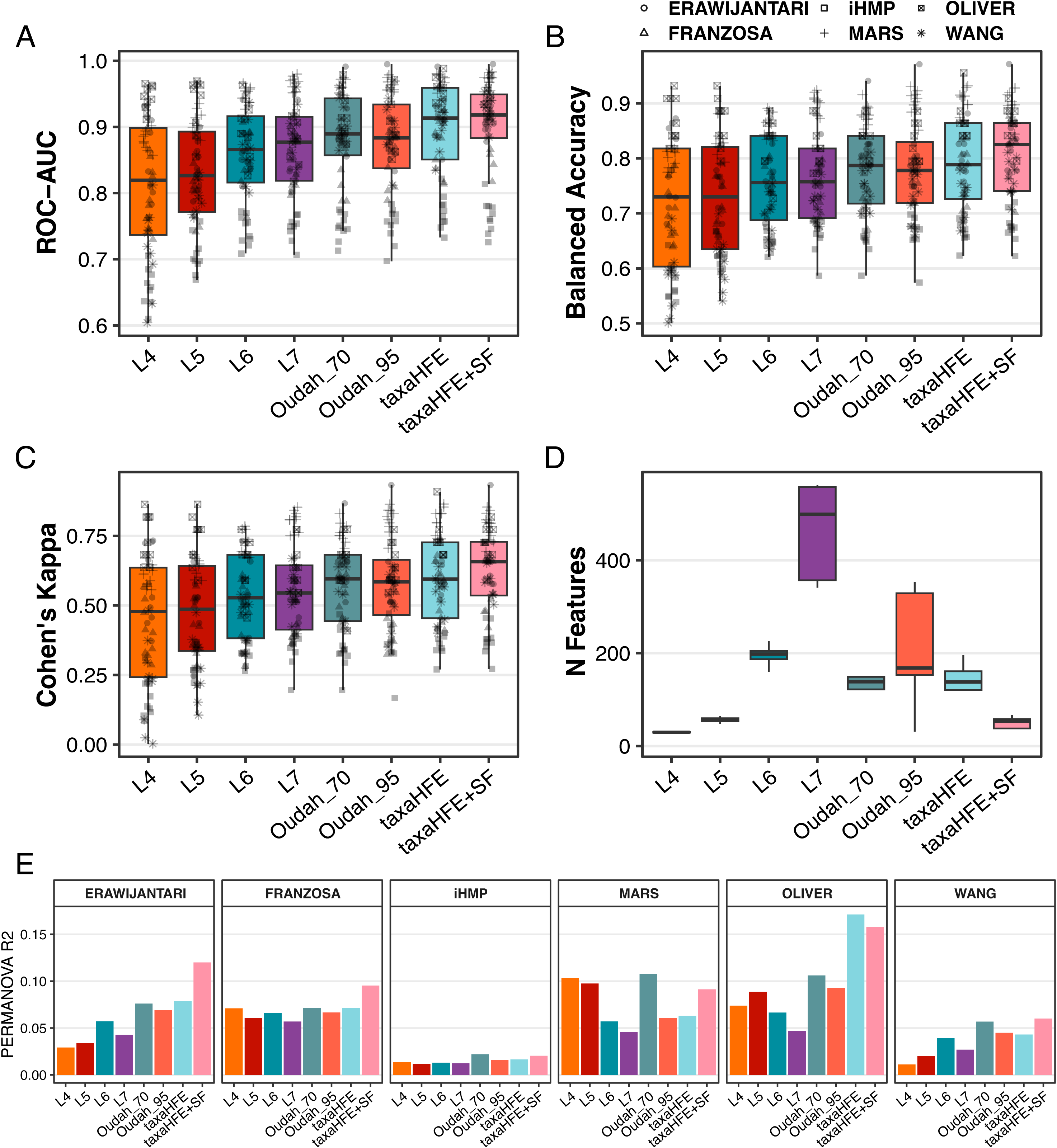
Comparisons made used summarized microbiome data (L4-L7, order through species), a previously published hierarchical feature engineering program (Oudah) employing two different internal correlation cutoffs (Pearson 0.7 and 0.95), and TaxaHFE with and without the super-filter. Machine learning performance metrics (A-C) and number of features used for model building (D). (E) Variance explained in PERMANOVA models by the composition of the features relative to the response variable used.

In addition to performance improvements, TaxaHFE (+SF) utilizes less features than comparable methods (Figure 2D). Models utilizing species-level features (after abundance and prevalence filters, see methods) utilized 469 (s.d. = 97) species on average, compared with the 45 (s.d. = 22) taxonomic features used by TaxaHFE (+SF) on average. Additionally, TaxaHFE (+SF) selects 95 less features on average compared to a previously published HFE algorithm by Oudah and Henschel (Oudah and Henschel, 2018). Importantly, TaxaHFE’s reduced feature set generally captures more variance in community composition compared to existing hierarchical feature engineering methods or summarizing to a specific taxonomic level (Figure 2E).

### TaxaHFE can reduce features for both categorical and continuous predictions

A previous effort towards hierarchical feature engineering produced an algorithm which maximized a performance metric with respect to a categorical variable (Oudah and Henschel, 2018). We designed TaxaHFE to handle both categorical and continuous variables. To illustrate this, we used normalized antibiotic resistance gene abundance as a continuous or categorical response variable for selecting taxa using previously published data (Oliver *et al*., 2022). Specifically, we only analyzed samples in the highest or lowest ARG abundance quartile. Using categorical ARG abundance (low quartile vs high quartile), TaxaHFE, with or without the default super-filter, selected 4 and 14 taxon features respectively. Using these features as input to a random forest classification model predicting categorical ARG abundance (high vs low) resulted in mean ROC-AUC values of 0.941 (+SF) and 0.952 (-SF) (Figure 3). Features summarized at the L4 level performed slightly better in classification models than TaxaHFE+SF (L4 (order) ROC-AUC: 0.943). The previously published HFE algorithm by Oudah and Henschel (Oudah and Henschel, 2018) selected 134 features, a considerable increase from TaxaHFE (+SF) 4 features. Even with far fewer features, TaxaHFE (+SF) achieved better performance metrics compared to models built with Oudah engineered features (ROC-AUC: TaxaHFE+SF, 0.941; Oudah_70, 0.915).

**Figure 3:**
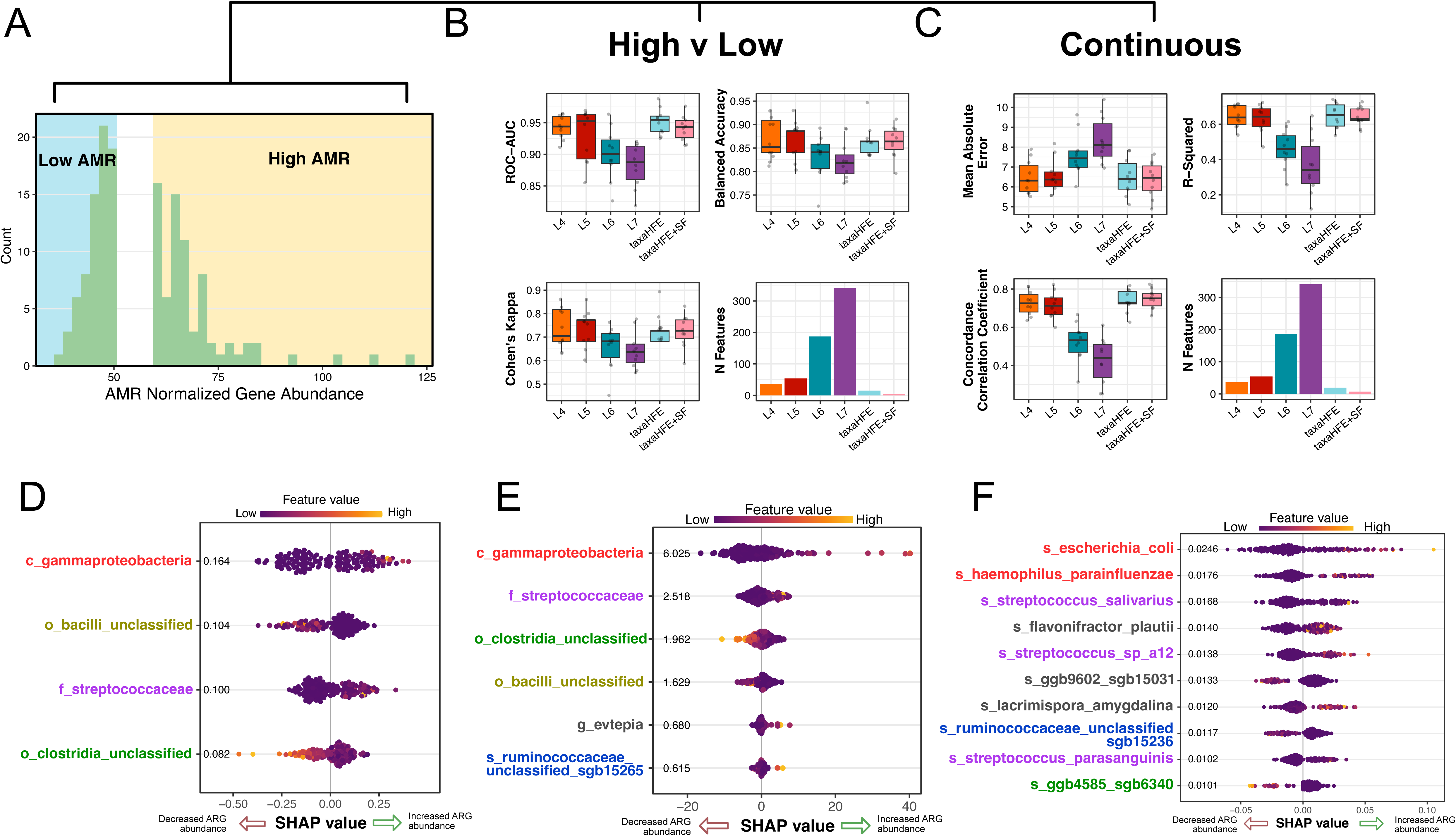
Analysis of TaxaHFE on categorical and continuous data. Histogram of antibiotic resistance gene abundance, showing just the top quartile and bottom quartile of the cohort (A). These data were analyzed as a categorical factor (low vs high ARG abundance) or continuous ARG abundance, and the (B, C) boxplots show the performance of machine learning models utilizing summarized taxonomic data or taxonomic hierarchical feature engineering. (D-F) Shapley values of the most informative features from a model built using TaxaHFE+SF data as input for both categorical and continuous ARG abundance are shown, as well as a model built only using L7 level data against the categorical ARG outcome. Colored font levels share a common taxonomic group with TaxaHFE.

When analyzing continuous ARG abundance, TaxaHFE, with or without the default super-filter, selected 6 and 18 taxon features respectively. Using these features as input to a random forest regression model predicting continuous ARG abundance resulted in mean R^2^ (coefficient of determination) values of 0.644 (+SF) and 0.649 (-SF) (Figure 3C). Like the categorical example, TaxaHFE+SF performed slightly behind the L4 model (by mean R^2^ performance), which used L4 (order) summarized data (0.645), yet a post-hoc test showed these differences were not significant, and the L4 model used far more features than TaxaHFE+SF (36 versus 6). No data is shown for the Oudah and Henschel algorithm as that algorithm does not support continuous outcome variables. In summary, TaxaHFE maximized performance while minimizing features for both categorical and continuous outcomes.

Both the categorical and continuous example resulted in models utilizing similar features (Figure 3D, E). Compared to a model built using Level 7 features (species), TaxaHFE identified higher taxonomic levels to discriminate categorical ARG abundance. These higher taxonomic levels, such as the class *Gammaproteobacteria* and family *Streptococcaceae,* resulted in nearly an order of magnitude higher SHAP values, suggesting TaxaHFE identified features that were more influential to model predictions than models built using species level data.

### TaxaHFE also works on hierarchically organized diet data

Since TaxaHFE works with hierarchical features, we sought to examine its utility beyond microbiome data. Dietary data represented as food trees could also be a useful feature set to apply hierarchical feature engineering. To test this, we utilized a previously published dietary food tree and tested TaxaHFE’s ability to select features which explain average fiber intake (Baldiviez *et al*., 2017). After prevalence and abundance filters, 456 foods were assessed. TaxaHFE+SF selected 4 features as the most informative for predicting average fiber intake, nearly a 99% reduction from the 456 features used as input. When assessed using the concordance correlation coefficient, a measure of both correlation and accuracy, Bonferroni corrected EMM post-hoc test revealed that TaxaHFE+SF performed significantly better than summarized features in a random forest (*p* < 0.05) (Figure 4A). We next used Shapley values to determine which features were most important to a model built with TaxaHFE+SF (Figure 4B) compared to a model built with Level 7 features (Figure 4C). The most important features identified by TaxaHFE+SF were “other fruits” (Level 2), which include high fiber staples such as berries and avocados, and “dry beans peas other legumes nuts and seeds” (Level 1) (Figure 4B). In contrast, the predictive model performance is lower for Level 7 (Figure 4A) and the SHAP values, which indicate the magnitude of importance, are also lower (Figure 4C) than those identified by TaxaHFE+SF (Figure 4B). In summary, the use of TaxaHFE improved both model performance and feature importance when used with dietary data arranged in food trees.

**Figure 4:**
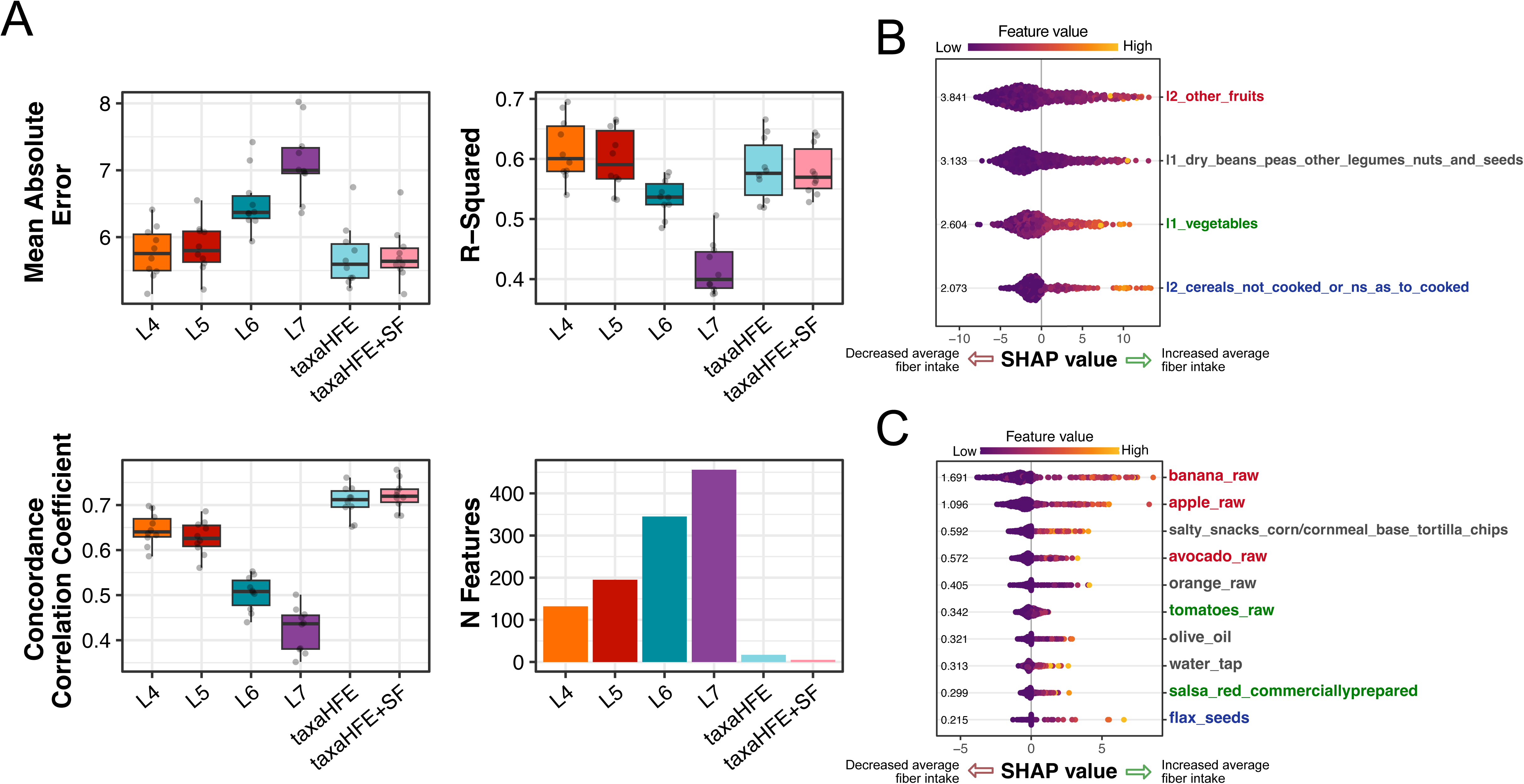
Testing TaxaHFE on hierarchically organized dietary data. Machine learning performance metrics of models predicting average fiber intake using food tree data (A) and number of features used for model building. Shapley values (B-C) of the most informative features from a model built using TaxaHFE+SF data as input or L6 as input. Colored font levels are those shared with TaxaHFE.

## DISCUSSION

A common practice in microbiome analyses is the summarization of abundance data at a single taxonomic level. In one review examining 419 microbiome studies, the authors found that the genus level was the most commonly summarized taxonomic level, followed by analysis at the phylum level (Kleine Bardenhorst *et al*., 2021). However, the discernable variation in microbial composition with respect to a trait or response is strongly dependent on the taxonomic level analyzed (Martiny *et al*., 2015). Thus, an analysis at a single taxonomic level, especially at a level which is not optimized for the trait or response of interest, could lead investigators to incorrectly conclude the absence of a relationship between microbiome and response. Our results support this; to explain ARG abundance, a model built using species level data would perform significantly worse than a model built using order level or TaxaHFE preprocessed data (Figure 3).

Here we propose an algorithm for the feature engineering of hierarchically organized data, particularly microbiome abundance data or dietary data represented using food trees. Currently, few algorithms exist to reduce the high dimensionality found in microbiome data with respect to the taxonomic structure inherent in microbial features. Oudah and Henschel (Oudah and Henschel, 2018) provided an excellent implementation; however, the Oudah algorithm is unable to handle: 1) a continuous response variable 2) non-bacterial features and 3) abundances that are not relative abundances. At the core of TaxaHFE is a random forest, which is a particularly capable ML algorithm for handling both continuous and categorical response variables. Moreover, there is no need for specific feature names; TaxaHFE bases its understanding of hierarchical levels purely based on the use of a separator (“|”) between levels. As such, any hierarchically organized data can be used as input.

One important shared logic of both the Oudah algorithm and our algorithm is the use of the parent taxon as the taxon to “beat” in the competitions. This decision was inspired by an early implementation of hierarchical feature engineering (Ristoski and Paulheim, 2014), for which the goal of the algorithm was to choose the most valuable features from the highest taxonomic levels possible. Indeed, when the goal is feature reduction, and the parent taxon and child taxa contain redundant information relative to a response of interest, choosing the parent taxon results in a feature that is likely representative of the child taxa in some way. The reverse is not necessarily true.

Other feature reduction programs such as Lefse (based on linear discriminant analysis) (Segata *et al*., 2011) and Boruta (based on random forests) (Kursa and Rudnicki, 2010) are often used, particularly with microbiome data. However, when all taxonomic levels are supplied to these programs, these programs often choose features that carry redundant abundance information. For example, when using Lefse to select features based the Erawijantari study, the output contained *Lactobacillales*, *Streptococcaceae*, and *Streptococcus* (Supplemental Figure 2A), which are all directly related features that carry nested but redundant feature abundance information (i.e., the abundance of *Streptococcus* is contained within the abundance of *Streptococcaceae*). And while Boruta performs slightly better than TaxaHFE in almost every case (Supplemental Figure 2B), it exhibits similar behavior as Lefse, choosing features with redundant feature abundance information (Supplemental Figure 2C). For some types of analyses this behavior is desirable. However, for explicit feature reduction we suggest that TaxaHFE’s method of removing overlapping features is usually preferable to aid interpretation of biological data.

Importantly, TaxaHFE preprocessed data improves the performance of machine learning models, especially compared to models produced using the lowest taxonomic levels available (e.g., species). This is perhaps not altogether surprising; indeed, a common observation across many microbiome studies is the highly personalized nature of microbiomes (Oliver *et al*., 2021). It stands to reason then, that while utilizing the most resolved taxonomic data might best highlight differences person to person, it will often mask more generalizable microbial responses within heterogeneous cohorts. Hierarchical feature engineering allows for the capture of these more generalizable responses, concomitantly increasing the accuracy of machine learning models in the process. We note, however, that if the goal is a generalizable model, using feature engineering inside of a cross-validation strategy is important to avoid data leakage (Aldehim and Wang, 2017). We briefly examined the similarity of features selected across k = 3-fold partitions of the data and found a high amount of dissimilarity (83%, data not shown) among the features selected in each fold. One reason for this variability is the small number of samples in the published studies we used to assess TaxaHFE (mean 262 samples). Like other feature reduction tools, we expect the performance of TaxaHFE will suffer when samples are limited.

Overall, perhaps the most important aspect of TaxaHFE is the gains in interpretability. In the microbiome example of the status quo method of using the lowest level taxa (Figure 3F), it is difficult to interpret the meaning of the ten species selected, each with a very low SHAP value. But with TaxaHFE (Figure 3D and 3E), it is readily apparent that the class *Gammaproteobacteria* and family *Streptococcaceae* are top predictors of antimicrobial resistance in the human gut microbiome and their SHAP values are an order of magnitude higher. The relationship between *Gammaproteobacteria* and antibiotic resistance has been shown previously in a Hi-C study linking ARGs to their microbial hosts (Stalder *et al*., 2019). Moreover, when considered together TaxaHFE selected features also appear to explain more compositional variance than using a single taxonomic level (Figure 2E). In the diet example (Figure 4C), the model based on the lowest taxonomic level is again difficult to interpret, with 456 individual food items (e.g., apples, bananas, avocados, oranges, olive oil, salty corn snacks, flax seeds, etc.) predictive of fiber intake. TaxaHFE reports low intake of other_fruits (non-citrus), legumes/nuts/seeds, uncooked cereals, and vegetables of low fiber intake. The results from TaxaHFE suggest, for example, that it is the entire class of legumes/nuts/seeds that is predictive, not just flax seeds, which is more reasonable. For this reason, dietary data is traditionally summarized and reported at the highest taxonomic level (e.g., fruits, vegetables, dairy, meat, etc.). However, nuances like the association of low intake of uncooked cereals with low fiber intake will be missed with such standard summary variables.

One shortcoming of TaxaHFE is its speed. While it is not a memory intensive program, it does rely on relatively inefficient for-loops to iterate over each taxon in each taxonomic level. For example, in a microbiome dataset with 4640 features, TaxaHFE took 2 min 44 sec, whereas the Oudah algorithm took only 13 seconds on a 2.3 GHz Quad-Core Intel® Core i7 machine. Future iterations of TaxaHFE could utilize a more distributed parallelization implementation.

Another limitation of TaxaHFE is true of all machine learning algorithms in that large high-quality data sets are needed. Without sufficient information in the data (i.e., no relationship between the response variable and the features) all ML algorithms will suffer. However, what may not be obvious to users is that in the absence of information, machine learning algorithms will still report results, which often represent noise. Methods for avoiding pitfalls of machine learning have been described elsewhere (Whalen *et al*., 2021), but we would be remiss to not echo them here. Specifically, we implore users to not merely trust models based on their accuracy alone, but to also investigate the features utilized for making predictions.

## CONCLUSION

We demonstrate that TaxaHFE dramatically reduces the feature space of hierarchically organized data while generally increasing the performance of downstream machine learning models and improving interpretability. While our examples come from microbiology and nutrition research, TaxaHFE could be used with any data set that has hierarchically related features. Moreover, TaxaHFE removes the prerequisite of choosing a taxonomic level that captures the most information relative to a response of interest, removing the need to ask, “At what taxonomic level should I analyze my data?”. Overall, we suggest that hierarchical feature engineering can lead to more accurate and interpretable models.

## ACKNOWLEDGEMENTS

We would like to acknowledge Elizabeth Chin for early TaxaHFE discussions and Stephanie Wilson and Sarah Blecksmith for useful feedback provided while code testing. We would like to thank Rachel Waymack and Jules Larke for thoughtful comments and edits. A.O. was supported by an ORISE fellowship administered by Oak Ridge Associated Universities through an interagency agreement between the U.S. Department of Energy and the U.S. Department of Agriculture, Agricultural Research Service.

## REFERENCES

Aldehim, G. and Wang, W. (2017) Determining appropriate approaches for using data in feature selection. Int. J. Mach. Learn. Cybern., 8, 915–928.

Baldiviez, L.M., et al. (2017) Design and implementation of a cross-sectional nutritional phenotyping study in healthy US adults. BMC Nutr., 3, 1–13.

Bellman, R.E. (2003) Dynamic Programming Courier Corporation.

Bevilacqua, S., et al. (2021) The use of taxonomic relationships among species in applied ecological research: Baseline, steps forward and future challenges.

Choi, Y., et al. (2022) A Guide to Dietary Pattern–Microbiome Data Integration. J. Nutr., 152, 1187–1199.

Erawijantari, P.P., et al. (2020) Influence of gastrectomy for gastric cancer treatment on faecal microbiome and metabolome profiles. Gut, 69, 1404–1415.

Franzosa, E.A., et al. (2019) Gut microbiome structure and metabolic activity in inflammatory bowel disease. Nat. Microbiol.

Glur, C., et al. (2020) data.tree: General Purpose Hierarchical Data Structure.

Greenwell, B. (2021) fastshap: Fast Approximate Shapley Value.

Hand, D.J. and Till, R.J. (2001) A Simple Generalisation of the Area Under the ROC Curve for Multiple Class Classification Problems. Mach. Learn., 45, 171–186.

Jacobs, D.R. and Steffen, L.M. (2003) Nutrients, foods, and dietary patterns as exposures in research: a framework for food synergy. Am. J. Clin. Nutr., 78, 508S–513S.

Johnson, A.J., et al. (2019) Daily Sampling Reveals Personalized Diet-Microbiome Associations in Humans. Cell Host Microbe, 25, 789–802.e5.

Kable, M.E., et al. (2022) Tree-Based Analysis of Dietary Diversity Captures Associations Between Fiber Intake and Gut Microbiota Composition in a Healthy US Adult Cohort. J. Nutr., 152, 779–788.

Kleine Bardenhorst, S., et al. (2021) Data Analysis Strategies for Microbiome Studies in Human Populations—a Systematic Review of Current Practice. mSystems.

Kuhn, M., et al. (2023) yardstick: Tidy Characterizations of Model Performance.

Kuhn, M. and Wickham, H. (2020) Tidymodels: a collection of packages for modeling and machine learning using tidyverse principles.

Kursa, M.B. and Rudnicki, W.R. (2010) Feature selection with the boruta package. J. Stat. Softw.

Lenth, R., et al. (2018) Emmeans. R Packag. version 1.15-15.

Lloyd-Price, J., et al. (2019) Multi-omics of the gut microbial ecosystem in inflammatory bowel diseases. Nature.

Mars, R.A.T., et al. (2020) Longitudinal Multi-omics Reveals Subset-Specific Mechanisms Underlying Irritable Bowel Syndrome. Cell.

Martiny, A.C., et al. (2012) Phylogenetic conservatism of functional traits in microorganisms. ISME J. 2013 74, 7, 830–838.

Martiny, J.B.H., et al. (2015) Microbiomes in light of traits: A phylogenetic perspective. Science (80-.)., 350.

Mayer, M. and Stando, A. (2023) shapviz: SHAP Visualizations.

Muller, E., et al. (2022) The gut microbiome-metabolome dataset collection: a curated resource for integrative meta-analysis. npj Biofilms Microbiomes.

Nembrini, S., et al. (2018) The revival of the Gini importance? Bioinformatics, 34, 3711–3718.

Oksanen, J. et al. (2019) vegan: Community Ecology Package. R package version 2.5-2. Cran R.

Oliver, A., et al. (2022) Association of Diet and Antimicrobial Resistance in Healthy U.S. Adults. MBio, 13.

Oliver, A., et al. (2021) High-Fiber, Whole-Food Dietary Intervention Alters the Human Gut Microbiome but Not Fecal Short-Chain Fatty Acids. mSystems, 6.

Oudah, M. and Henschel, A. (2018) Taxonomy-aware feature engineering for microbiome classification. BMC Bioinformatics, 19, 1–13.

Pinheiro, J., et al. (2019) nlme: Linear and Nonlinear Mixed Effects Models.

Ristoski, P. and Paulheim, H. (2014) Feature Selection in Hierarchical Feature Spaces. In, Lecture Notes in Computer Science (including subseries Lecture Notes in Artificial Intelligence and Lecture Notes in Bioinformatics)., pp. 288–300.

Segata, N., et al. (2011) Metagenomic biomarker discovery and explanation. Genome Biol., 12, R60.

Stalder, T., et al. (2019) Linking the resistome and plasmidome to the microbiome. ISME J. 2019 1310, 13, 2437–2446.

Štrumbelj, E. and Kononenko, I. (2014) Explaining prediction models and individual predictions with feature contributions. Knowl. Inf. Syst.

Wang, X., et al. (2020) Aberrant gut microbiota alters host metabolome and impacts renal failure in humans and rodents. Gut.

Wetterstrand., K.A. DNA Sequencing Costs: Data from the NHGRI Genome Sequencing Program (GSP).

Whalen, S., et al. (2021) Navigating the pitfalls of applying machine learning in genomics. Nat. Rev. Genet. 2021 233, 23, 169–181.

Wright, M.N. and Ziegler, A. (2017) rangerL: A Fast Implementation of Random Forests for High Dimensional Data in C++ and R. J. Stat. Softw., 77.

