## Supplementary figures and images for "TaxaHFE: A machine learning approach to collapse microbiome datasets using taxonomic structure"

### Supplemental Figure 1

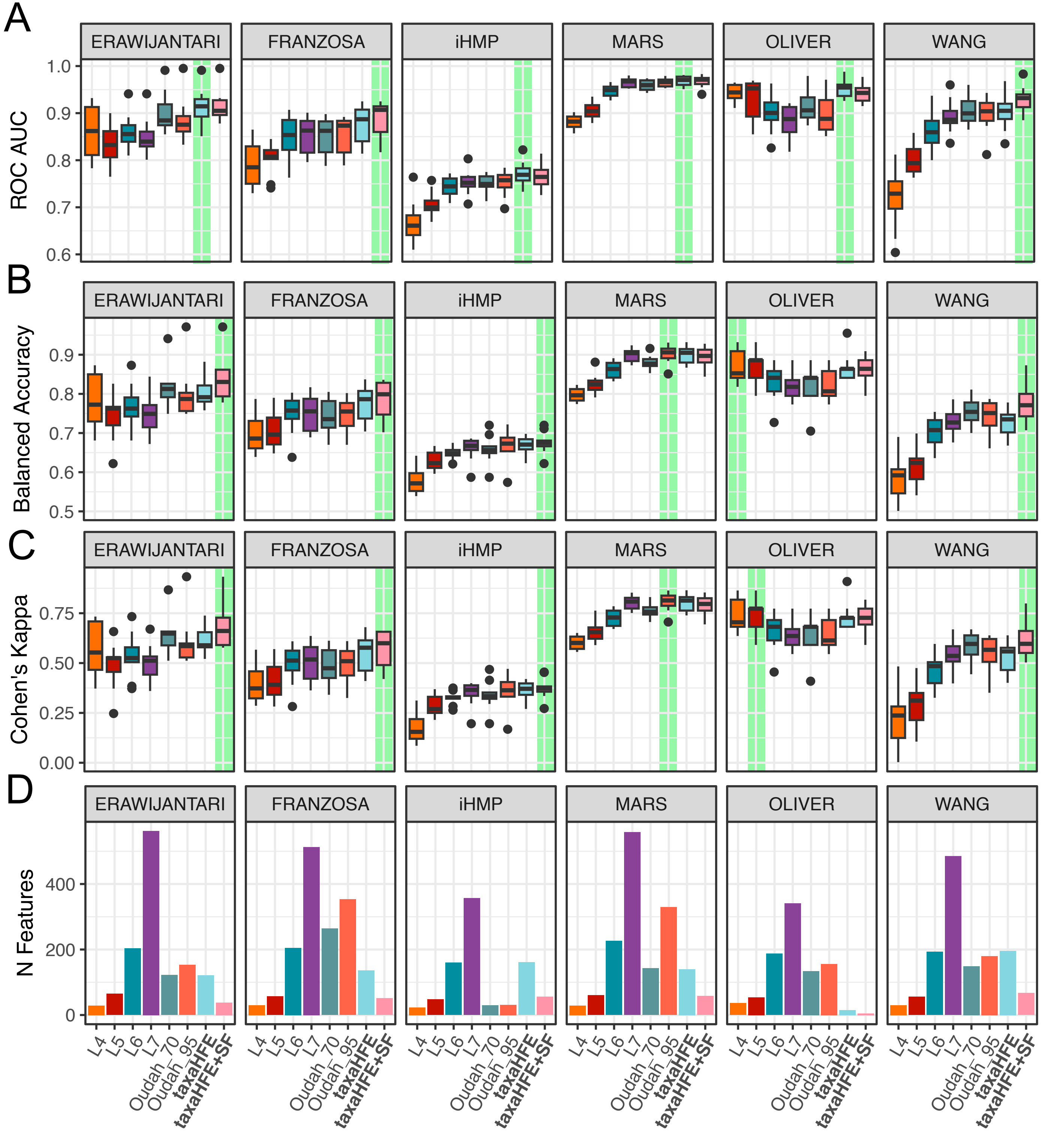
