## Supplemental Figure 2 for "TaxaHFE: A machine learning approach to collapse microbiome datasets using taxonomic structure"

A

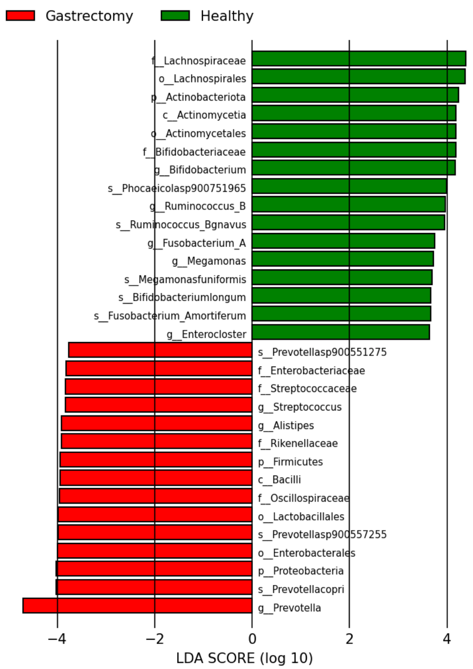


B

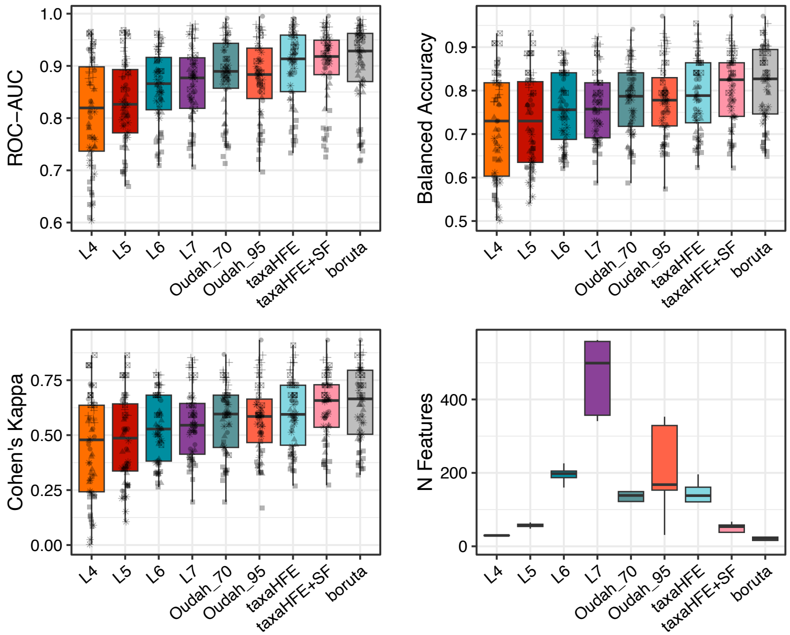


C

### Selected features for Oliver et al., 2022

k_bacteria_p_firmicutes_c_clostridia_o_clostridia_unclassified
k_bacteria_p_firmicutes_c_clostridia_o_clostridia_unclassified_f_clostridia_unclassified
k_bacteria_p_firmicutes_c_clostridia_o_clostridia_unclassified_f_clostridia_unclassified_g_ggb4585
k_bacteria_p_firmicutes_c_clostridia_o_clostridia_unclassified_f_clostridia_unclassified_g_ggb4585_s_ggb4585_sgb6340
k_bacteria_p_firmicutes_c_clostridia_o_clostridia_unclassified_f_clostridia_unclassified_g_ggb4585_s_ggb4585_sgb6340_t_sgb6340
k_bacteria_p_firmicutes_c_bacilli_o_lactobacillales
k_bacteria_p_firmicutes_c_bacilli_o_lactobacillales_f_streptococcaceae
k_bacteria_p_firmicutes_c_bacilli_o_lactobacillales_f_streptococcaceae_g_streptococcus
k_bacteria_p_firmicutes_c_bacilli_o_bacilli_unclassified
k_bacteria_p_firmicutes_c_bacilli_o_bacilli_unclassified_f_bacilli_unclassified
k_bacteria_p_firmicutes_c_bacilli_o_bacilli_unclassified_f_bacilli_unclassified_g_bacilli_unclassified
k_bacteria_p_actinobacteria_c_actinobacteria_o_actinomycetales_f_actinomycetaceae_g_actinomyces
k_bacteria_p_actinobacteria_c_actinobacteria_o_actinomycetales_f_actinomycetaceae_g_actinomyces_s_actinomyces_graevenitzii_t_sgb17130
k_bacteria_p_proteobacteria_c_deltaproteobacteria_o_desulfovibrionales_f_desulfovibrionaceae_g_desulfovibrio
k_bacteria_p_proteobacteria_c_gammaproteobacteria
k_bacteria_p_proteobacteria_c_gammaproteobacteria_o_enterobacterales
k_bacteria_p_proteobacteria_c_gammaproteobacteria_o_enterobacterales_f_enterobacteriaceae
k_bacteria_p_proteobacteria_c_gammaproteobacteria_o_enterobacterales_f_enterobacteriaceae_g_escherichia
k_bacteria_p_proteobacteria_c_gammaproteobacteria_o_enterobacterales_f_enterobacteriaceae_g_escherichia_s_escherichia_coli
k_bacteria_p_proteobacteria_c_gammaproteobacteria_o_enterobacterales_f_enterobacteriaceae_g_escherichia_s_escherichia_coli_t_sgb10068
k_bacteria_p_proteobacteria_c_gammaproteobacteria_o_pasteurellales
k_bacteria_p_proteobacteria_c_gammaproteobacteria_o_pasteurellales_f_pasteurellaceae
k_bacteria_p_proteobacteria_c_gammaproteobacteria_o_pasteurellales_f_pasteurellaceae_g_haemophilus
k_bacteria_p_proteobacteria_c_gammaproteobacteria_o_pasteurellales_f_pasteurellaceae_g_haemophilus_s_haemophilus_parainfluenzae
k_bacteria_p_proteobacteria_c_gammaproteobacteria_o_pasteurellales_f_pasteurellaceae_g_haemophilus_s_haemophilus_parainfluenzae_t_sgb9712_group
